# Fast Approximate IsoRank for Scalable Global Alignment of Biological Networks

**DOI:** 10.1101/2023.03.13.532445

**Authors:** Kapil Devkota, Anselm Blumer, Lenore Cowen, Xiaozhe Hu

## Abstract

A well-studied approximate version of the graph matching problem is directly relevant for the study of protein-protein interaction networks. Called by the computational biology community Global Network Alignment, the two networks to be matched are derived from the protein-protein interaction (PPI) networks from organisms of two different species. If these two species evolved recently from a common ancestor, we can view the two PPI networks as a single network that evolved over time. It is the two versions of this network that we want to align using approximate graph matching. The first spectral method for the PPI global alignment problem proposed by the biological community was the IsoRank method of Singh et al. This method for global biological network alignment is still used today. However, with the advent of many more experiments, the size of the networks available to match has grown considerably, making running IsoRank unfeasible on these networks without access to state of the art computational resources. In this paper, we develop a new IsoRank approximation, which exploits the mathematical properties of IsoRank’s linear system to solve the problem in quadratic time with respect to the maximum size of the two PPI networks. We further propose a computationally cheaper refinement to this initial approximation so that the updated result is even closer to the original IsoRank formulation. In experiments on synthetic and real PPI networks, we find that the results of our approximate IsoRank are not only nearly as accurate as the original IsoRank results but are also much faster, which makes the global alignment of large-scale biological networks feasible and scalable.

## 1 Introduction

As has been noted in the social network and pattern recognition applications communities as well [6, 11], spectral methods have enjoyed some success in many variants of the approximate graph matching problem, including for the *Global Network Alignment* problem studied by computational biologists for aligning protein-protein interaction (PPI) networks from different species. The first spectral method for the PPI global alignment problem proposed by the biological community was the IsoRank method of Singh et al. [16]. This method for global biological network alignment is still used today. With the advent of many more experiments, the size of the networks available to match has grown considerably making running IsoRank unfeasible on these networks without access to state of the art computational resources. In this paper, we develop a new IsoRank approximation that exploits the eigenproperties of IsoRank’s linear system to solve the problem in quadratic time with respect to the maximum size of the two PPI networks. We further propose a computationally cheaper refinement to this initial approximation so that the updated result is even closer to the original IsoRank formulation.

We demonstrate the quality of our approximation in two different settings. First, in synthetic experiments, we create random graphs using the Erdős Rényi [7] and Barabási-Albert models [1], and ask IsoRank to recover the graph isomorphism between our graphs and a random node permutation. We add various levels of noise to the node similarities (which are either 0 or 1 indicating the random node permulation). We measure the quality of our IsoRank approximation for three ranges of the noise level: a low-error regime where IsoRank itself will recover the matching of the true isomorphism, an intermediate *E* where IsoRank only gets 80%-90% of the true matching, and a noisy E where IsoRank gets only 70% - 75% of the true matching. In each case, we show how close approximate IsoRank comes, as well as demonstrating the considerable savings in CPU time. Second, we return to the biological domain, and consider real-world networks from 5 species: mouse, rat, fly, yeast and human. In this case, the more distant species can be seen as having a noisier E value, since the sequence similarity diverges over evolutionary time (i.e. it is much easier to align mouse and rat than human and fly). However, in addition to an increasing *E* distance of node similarity (measured by the Biology industry standard algorithm, BLAST [2]), we don’t have the same number of nodes in each PPI network, we have inserted and deleted edges in each PPI network, and finally it is assumed we have only a noisy and incomplete sample of the true PPI networks. Because we of course do not have ground truth in this setting, we instead adopt the popular measures of quality that are common in the bioinformatics community for the Global Network Alignment problem, namely the EC, LCCS measures [8] of structural alignment quality, as well as the AFS measure [8] which measures how often aligned proteins are known to be involved in similar biological functions across the two species (see Section 3.2 below for definitions). Additional alternative ways of measuring quality of biological network alignment appear in [17, 14, 13, 5]

## 2 Algorithm

### 2.1 IsoRank

Let us first recall the IsoRank algorithm. The key observation is that IsoRank looks at the *tensor product* of two graphs, which is essential for developing our approximation algorithm. The relation to tensor product was also previously observed by Zhang et al. [10, 18, 19].

Consider two simply connected, undirected, and possibly weighted graphs 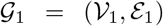 and 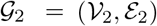. Their adjacency matrices are *A*_1_ and *A*_2_, respectively, and weighted degree matrices are *D*_1_:= diag(d_1_), 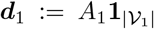, and *D*_2_:= diag(***d***_2_), 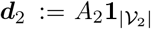, respectively. Thus, the corresponding transition matrices are

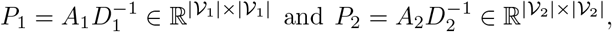

which are both column stochastic, i.e., 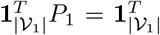 and 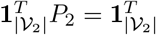

Next we consider the tensor product of the two graphs, which is a graph with 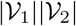 nodes and its adjacency matrix is defined as

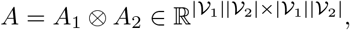

where ⊗ denotes the Kronecker product. Similarly, the transition matrix of the tensor product graph is

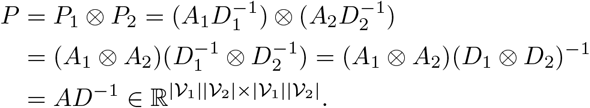

where 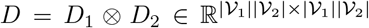 is the degree matrix of the tensor product graph. This is because

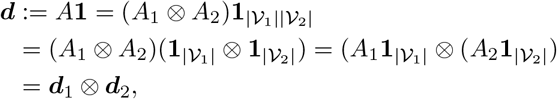

and

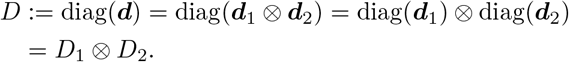

As suggested in [16], the basic version of IsoRank tries to find 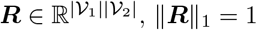, such that

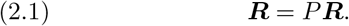

This is done by the power method, which is the following iterative procedure, for *k* = 0, 1, 2,…

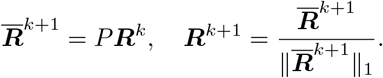

A modified version of IsoRank tries to find **R**, ||**R**||_1_ = 1, such that, for 0 ≤ *α* ≤ 1

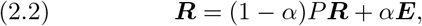

where 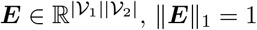, ||*E*||_1_ = 1 which typically contains domain specific information to improve the graph alignment result. For example, in the original IsoRank paper [16], besides the two PPI networks of the species, additional sequence similarity data, which is presented by 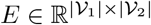, is required in order to perform the graph matching. Here *E_ij_* denotes how similar the amino acid sequence of the *i^th^* protein of 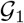 is to that of the *j^th^* protein of 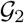. This sequence similarity information is obtained using the standard BLAST sequence alignment [2]. Since we use the tensor product notation in our paper, we have 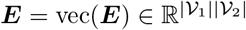, where vec(·) converts a matrix to a vector.

As suggested in [16], (2.2) is again solved iteratively as follows, for *k* = 0, 1, 2,…

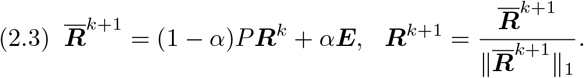

Other approximation methods can also be used to solve (2.2), e.g., [10].

#### Remark 1.

*Note that, in both iterative procedures, the normalization step is not needed if we have **R***^0^ ≥ 0, ||***R***^0^||_1_ = 1, ***E*** ≥ 0, *and* ||***E***||_1_ = 1. *This is because*

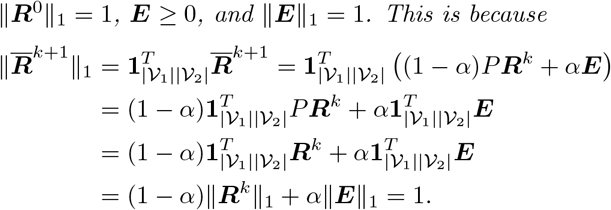

### 2.2 Approximate IsoRank

Let us first consider the basic version of IsoRank (2.1) and the key here is to notice that IsoRank is actually working on a tensor product graph of the original two graphs. Therefore, based on the Perron–Frobenius theorem, the basic IsoRank (2.1) tries to find the right eigenvector of the transition matrix *P* that corresponds to the unique eigenvalue 1. Therefore, ***R*** is nothing but the steady state distribution (SSD) ***π*** of the tensor product graph, which can be computed directly without using the power method at all since IsoRank works on a tensor product graph. Using the properties of the transition matrix *P* of a simply connected, undirected, and possibly weighted graph, the SSD can be obtained without any iterative procedure as follows,

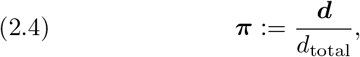

where 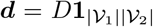 and 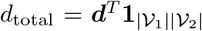 (which is the weighted total degree). This means, for the basic version IsoRank (2.1), we have ***R***:= ***π*** directly and there is no need of the power method or any iterative procedure which involves matrix vector multiplications.

Now let us consider the modified version of IsoRank (2.2), which is preferred in practice due to the incorporation of the extra information. Note that when *α* = 0, (2.2) reduces to (2.1) and, therefore, ***R*** = ***π***. On the other hand, when *α* = 1, we have ***R*** = ***E***. Thus, for 0 ≤ *α* ≤ 1, we can use the following convex combination to get an approximation

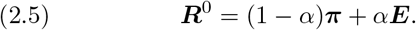

Again, there is no iterative procedure and no matrix-vector multiplications at all. The reason we call this approximation ***R***^0^ is that it can be used as an approximation directly, or as the initial guess of an iterative procedure for solving (2.2).

According to (2.2) and (2.5), we have

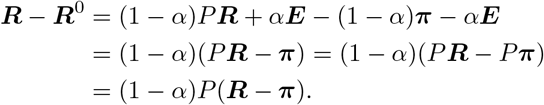

Let us introduce 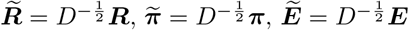, and 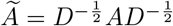. Then we have

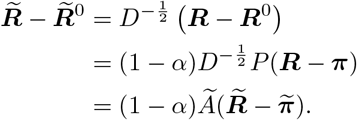

On the other hand, multiplying 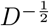 on the both sides of (2.2), we obtain

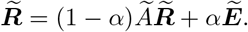

This implies

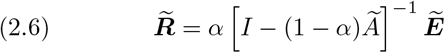

Since 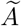 is symmetric and doubly stochastic, its eigenvalues are 1 = *λ*_1_ > *λ*_2_ ≥ *λ*_3_ ≥… ≥ *λ_n_*, 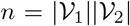. The corresponding eigenvectors are denoted by ***v**_i_*, *i* = 1,…, *n*, with ||***v**_i_*|| = 1 and 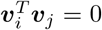, *i* ≠ *j*. It is easy to check that 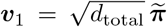. Since 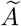 is symmetric, ***v**_i_*, *i* = 1,…, *n*, form an orthonormal basis and, thus 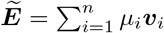 with 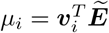. Note that,

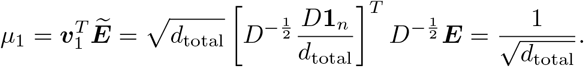

Now we are ready to present the following theorem which measures the approximation error of ***R***^0^.

#### Theorem 2.1.

*Let **R***^0^ *be defined as* (2.5), *for* 0 < *α* ≤ 1, *we have*

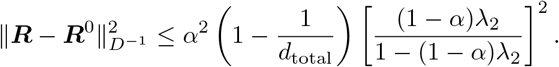

*where* 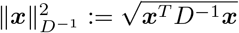.

*Proof.* Based on (2.6), for 0 < *α* ≤ 1,

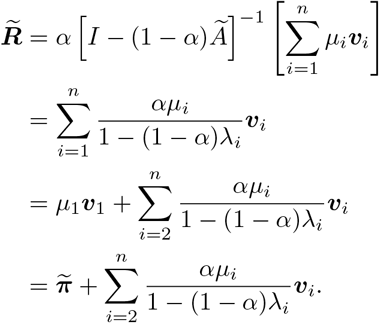

This means,

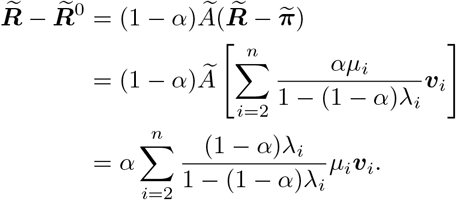

Therefore,

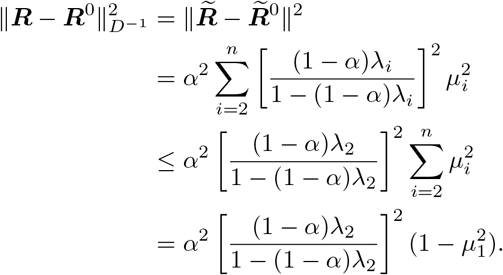

This completes the proof.

As we mentioned, if we are not satisfied with *R*^0^ (2.5), we can use it as an initial guess and perform power iterations. In practice, since it provide a good initial guess, we found that usually only one step of power iteration is needed which provides another approximation **R**^1^ as follows,

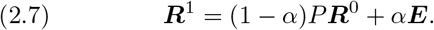

Note that, due to the fact that *P* = *P*_1_ ⊗ *P*_2_, the matrixvector multiplication step *y* = *P_x_* can be computed without explicitly forming *P*. We can use the following equivalent expression to compute *y*

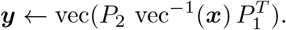

where vec^-1^(·) is the inverse operation of vec(·).

### 2.3 Computational Complexity

As we can see, the computational cost of using the iterative method (2.3) to solve IsoRank (2.2), Our approximation (2.5), and (2.7) mainly depends on the matrix-vector multiplication for computing *P**x*** in the iteration (if the implementation uses the tensor product, then this step becomes matrix-matrix multiplication) and vector addition for computing *ax* + *by*, 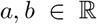. Let us assume the complexity for matrix-vector multiplication (or matrix-matrix multiplication in the tensor product implementation) is *t*_1_ and the complexity for vector addition is *t*_2_. Then the complexity of using the iterative method (2.3) to solve IsoRank (2.2) is *k*(*t*_1_ + *t*_2_), where *k* is the number of iterations used in the implementation. Using the standard convergence analysis of (2.3), at least 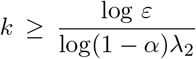 steps are requires to reach a given relative error tolerance *ε* (measured in || · ||_D-1_ norm). Here *λ*_2_ is the second largest eigenvalue of *P*, which happens to be the second largest eigenvalue of 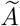 since *P* and 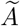 are similar.

For our approximation ***R***^0^, since we only use vector addition once, the complexity is *t*_2_. Finally, the approximation ***R***^1^ does one step of iteration using ***R***^0^ as the initial guess. Thus, its computational cost is *t*_2_ + (*t*_1_ + *t*_2_) = *t*_1_ + 2*t*_2_.

As we can see, our approximation’s main computational advantage is that there is no need for iterations. Consider the worst case that *λ*_2_ ≈ 1, let us choose *α* = 0.6 as used in our experiments. The iterative method (2.3) needs about *k* = 30 iterations to achieve *ε* = 10^-12^. This implies that our approximations are about 30 times faster than iterating to convergence here. Of course, the computational saving will depend on specific implementations and computers.

## 3 Experiments

### 3.1 Synthetic Networks

Our first set of experiments uses the Erdős Rényi (ER) random graphs [7] and the Barabási Albert (BA) random graphs [1]. For the ER graphs, to ensure the resulting random graphs are simply connected, we set the probability of using an edge to connect any two nodes to be 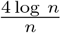. For the BA graphs, we use a tree graph consisting of 5 nodes and edge set {(1, 2), (1, 5), (2, 4), (3, 4)} as the seed graph. Each new node is connected to 5 existing nodes with a probability proportional to the number of edges the existing nodes already have.

To apply and test the IsoRank algorithm (2.2) and our two approximation algorithms, i.e., (2.5) and (2.7), we first generate a simply-connected random graph and then randomly permute the nodes to obtain another random graph. As suggested in [16], we choose *α* = 0.6 in (2.2) and all the tests. In addition, we generate different ***E*** by adding a different level of random noise to the edge weights of the correct random permutation. Starting with the correct random permutation, whose values are either 0 or 1, for a “good” ***E***, we add random noise between [-0.5, 0.5]. For a “bad” ***E***, we add random noise between [-0.7, 0.7]. When we add random noise between [-0.6, 0.6], we refer to the resulting ***E*** as “intermediate”. Once the random noise is added, we take absolute value component-wise and then normalize to ensure that ||**E**||_1_ = 1 and all the components are non-negative. Intuitively, the “good” ***E*** provides more information about the correct alignment, resulting in better accuracy than other choices of **E**. Finally, since the IsoRank(2.2) is solved by an iterative procedure, we stop the iteration when the Frobenius norm of the difference between two successive iterations is less than 10^-12^ or the maximal number of iterations, which is set to be 100, is reached. In all our experiments, it took less than 20 iterations to converge.

We report the accuracy, CPU time, ||·|| error, and ||·||_*D*-1_ error. The accuracy of the alignment is measured in terms of how many pairs of nodes between the original network and permuted network are predicted correctly (in percentage). Since we implement all the algorithms in Matlab, the CPU time is measured using the Matlab build-in functions `tic` and `toc`. Finally, we use the standard ||·|| norm to measure the error between the IsoRank result and its approximations. To verify the theory, we also look at the error using the weighted ||·||_*D*-1_ norm. The numerical results are reported in Table 1 through 6.

**Table 1:**
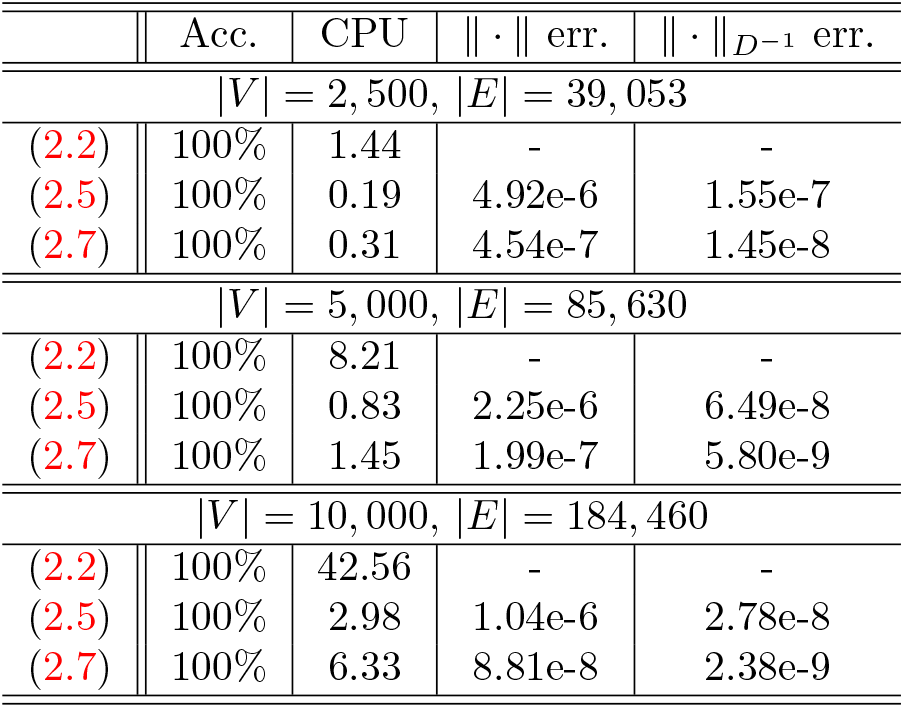
Erdős Rényi Random Graph: good ***E***

| | Acc. | CPU | $\ \cdot\ $ err. | $\ \cdot\ _{D^{-1}}$ err. |
| --- | --- | --- | --- | --- |
| $\vert V\vert = 2,500, \vert E\vert = 39,053$ | | | | |
| (2.2) | 100% | 1.44 | - | - |
| (2.5) | 100% | 0.19 | 4.92e-6 | 1.55e-7 |
| (2.7) | 100% | 0.31 | 4.54e-7 | 1.45e-8 |
| $\vert V\vert = 5,000, \vert E\vert = 85,630$ | | | | |
| (2.2) | 100% | 8.21 | - | - |
| (2.5) | 100% | 0.83 | 2.25e-6 | 6.49e-8 |
| (2.7) | 100% | 1.45 | 1.99e-7 | 5.80e-9 |
| $\vert V\vert = 10,000, \vert E\vert = 184,460$ | | | | |
| (2.2) | 100% | 42.56 | - | - |
| (2.5) | 100% | 2.98 | 1.04e-6 | 2.78e-8 |
| (2.7) | 100% | 6.33 | 8.81e-8 | 2.38e-9 |

**Table 2:** Erdős Rényi Random Graph: intermediate ***E***

| | Acc. | CPU | $\ \cdot\ $ err. | $\ \cdot\ _{D^{-1}}$ err. |
| --- | --- | --- | --- | --- |
| $\vert V\vert = 2,500, \vert E\vert = 38,802$ | | | | |
| (2.2) | 83.9% | 1.41 | - | - |
| (2.5) | 83.3% | 0.19 | 4.90e-6 | 1.56e-7 |
| (2.7) | 83.8% | 0.29 | 4.58e-7 | 1.46e-8 |
| $\vert V\vert = 5,000, \vert E\vert = 84,850$ | | | | |
| (2.2) | 83.6% | 7.88 | - | - |
| (2.5) | 83.0% | 0.83 | 2.31e-6 | 6.70e-8 |
| (2.7) | 83.6% | 1.43 | 2.08e-7 | 6.08e-9 |
| $\vert V\vert = 10,000, \vert E\vert = 183,929$ | | | | |
| (2.2) | 83.1% | 45.66 | - | - |
| (2.5) | 82.4% | 3.27 | 1.05e-6 | 2.82e-8 |
| (2.7) | 83.1% | 6.62 | 8.97e-8 | 2.43e-9 |

**Table 3:**
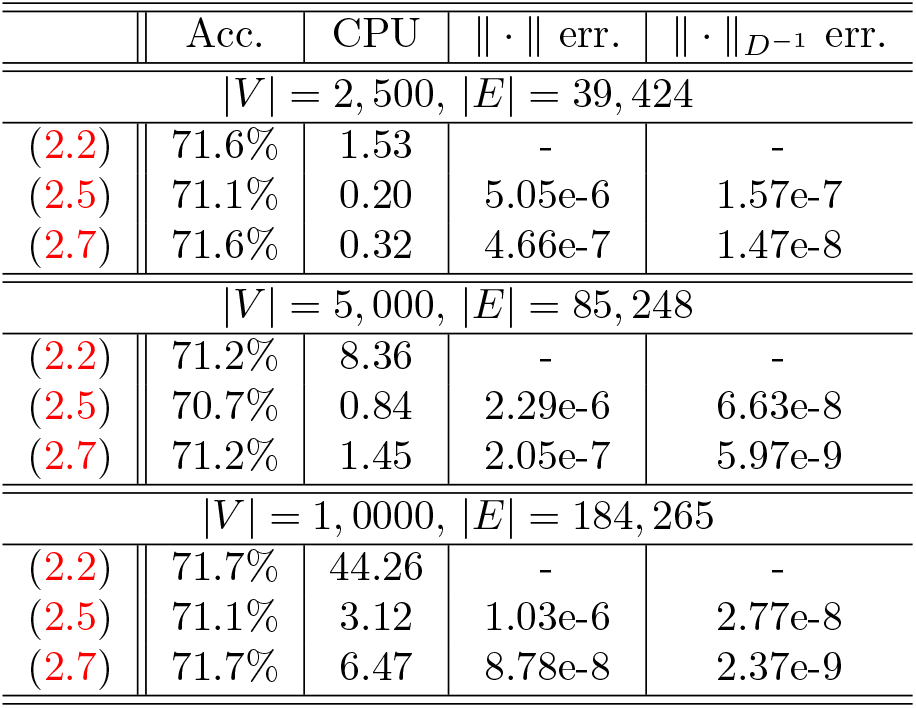
Erdős Rényi Random Graph: bad ***E***

| | Acc. | CPU | $\ \cdot\ $ err. | $\ \cdot\ _{D^{-1}}$ err. |
| --- | --- | --- | --- | --- |
| $\vert V\vert = 2,500, \vert E\vert = 39,424$ | | | | |
| (2.2) | 71.6% | 1.53 | - | - |
| (2.5) | 71.1% | 0.20 | 5.05e-6 | 1.57e-7 |
| (2.7) | 71.6% | 0.32 | 4.66e-7 | 1.47e-8 |
| $\vert V\vert = 5,000, \vert E\vert = 85,248$ | | | | |
| (2.2) | 71.2% | 8.36 | - | - |
| (2.5) | 70.7% | 0.84 | 2.29e-6 | 6.63e-8 |
| (2.7) | 71.2% | 1.45 | 2.05e-7 | 5.97e-9 |
| $\vert V\vert = 1,0000, \vert E\vert = 184,265$ | | | | |
| (2.2) | 71.7% | 44.26 | - | - |
| (2.5) | 71.1% | 3.12 | 1.03e-6 | 2.77e-8 |
| (2.7) | 71.7% | 6.47 | 8.78e-8 | 2.37e-9 |

**Table 4:** Barabaási Albert Random Graph: good ***E***

| | Acc. | CPU | $\ \cdot\ $ err. | $\ \cdot\ _{D^{-1}}$ err. |
| --- | --- | --- | --- | --- |
| $\vert V\vert = 2,500, \vert E\vert = 12,473$ | | | | |
| (2.2) | 100% | 3.37 | - | - |
| (2.5) | 100% | 0.21 | 4.69e-5 | 2.66e-6 |
| (2.7) | 100% | 0.38 | 1.19e-5 | 4.28e-7 |
| $\vert V\vert = 5,000, \vert E\vert = 24,975$ | | | | |
| (2.2) | 100% | 13.81 | - | - |
| (2.5) | 100% | 0.84 | 2.41e-5 | 1.32e-6 |
| (2.7) | 100% | 1.57 | 6.77e-6 | 2.10e-7 |
| $\vert V\vert = 10,000, \vert E\vert = 49,974$ | | | | |
| (2.2) | 100% | 38.85 | - | - |
| (2.5) | 100% | 3.15 | 1.21e-5 | 6.60e-7 |
| (2.7) | 100% | 5.18 | 3.62e-6 | 1.05e-7 |

**Table 5:** Barabáasi Albert Random Graph: intermediate ***E***

| | Acc. | CPU | $\ \cdot\ $ err. | $\ \cdot\ _{D^{-1}}$ err. |
| --- | --- | --- | --- | --- |
| $\vert V\vert = 2,500, \vert E\vert = 12,465$ | | | | |
| (2.2) | 87.3% | 3.22 | - | - |
| (2.5) | 84.9% | 0.19 | 4.41e-5 | 2.57e-6 |
| (2.7) | 87.2% | 0.35 | 1.14e-5 | 4.04e-7 |
| $\vert V\vert = 5,000, \vert E\vert = 24,976$ | | | | |
| (2.2) | 86.9% | 14.03 | - | - |
| (2.5) | 83.9% | 0.83 | 2.37e-5 | 1.33e-6 |
| (2.7) | 86.8% | 1.57 | 6.39e-6 | 2.11e-7 |
| $\vert V\vert = 10,000, \vert E\vert = 49,972$ | | | | |
| (2.2) | 86.6% | 37.43 | - | - |
| (2.5) | 84.1% | 3.08 | 1.22e-5 | 6.55e-7 |
| (2.7) | 86.5% | 5.11 | 3.48e-6 | 1.04e-7 |

**Table 6:**
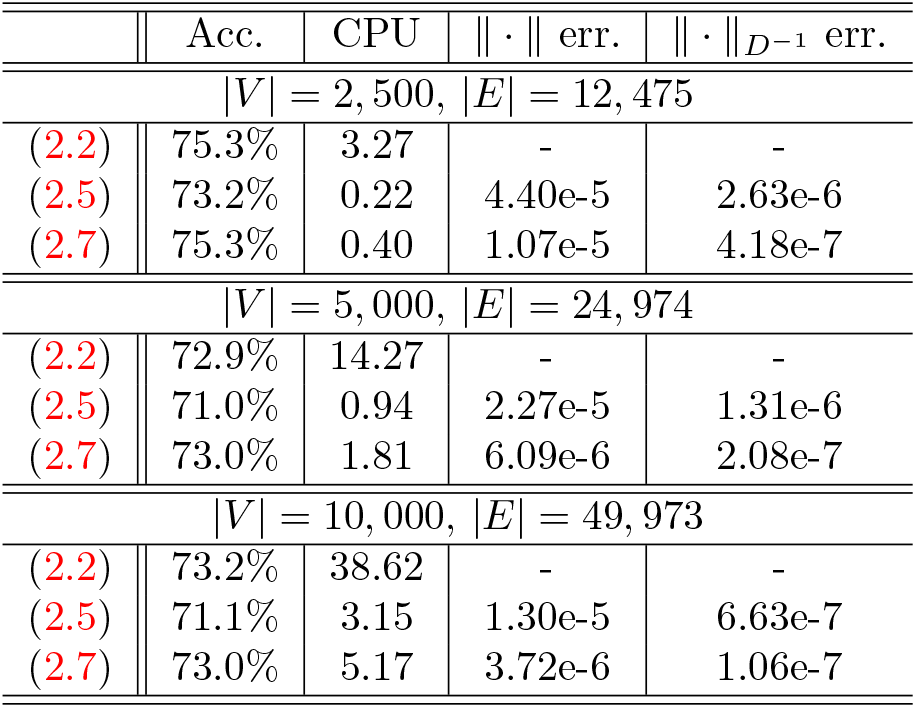
Barabási Albert Random Graph: bad ***E***

### 3.2 Biological Networks

We also tested the IsoRank approximations on real PPI networks. We extracted networks from the IntAct database [9]; these networks are constructed using all physical interactions detected using Coimmunoprecipitation (Co-IP) [3] and Yeast 2-Hybrid (Y2H) [12] experiments. We generated the networks for five species: *S* = {Human, Mouse, Rat, Fly, Baker’s Yeast}, where Human = *Homo sapiens*, Mouse = *Mus musculus*, Rat = Rattus norvegicus, Fly = *Drosophila melanogaster* and Baker’s yeast = *Saccharomyces cerevisiae.* The graph properties of these constructed PPI networks are provided in Table 7.

**Table 7:** Graph properties of IntAct networks. Table includes information regarding the number of nodes, number of edge, average node degree and average clustering coefficient for human, fly, mouse, rat and yeast networks.

| network | #nodes | #edges | avg degree | avg cc |
| --- | --- | --- | --- | --- |
| human | 32431 | 301529 | 18.595 | 0.069 |
| fly | 11247 | 42555 | 7.567 | 0.027 |
| mouse | 13967 | 36541 | 5.232 | 0.051 |
| rat | 10792 | 23315 | 4.320 | 0.075 |
| yeast | 6478 | 70638 | 21.808 | 0.285 |

From these 5 species, we can generate 10 distinct pairwise combinations of species, and for each pair of corresponding PPI networks, we performed experiments using the true IsoRank similarity matrix *R*, and its two approximations *R*^0^ and *R*^1^. Finally, for each obtained similarity matrix *R, R*^0^ and *R*^1^, we used the greedy matching algorithm to find the top 2000 protein mappings *M, M*_0_ and *M*_1_ respectively.

Additionally, to measure the biological properties of the mappings obtained from IsoRank and its approximations, we used three metrics: Edge Correctness (EC), Largest Common Connected Subgraph (LCCS) and Average Functional Similarity (AFS) [8]. A detailed description of these metrics is provided below.

#### Edge Correctness (EC)

Let 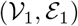 and 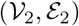 represent two networks, with 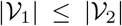 and let 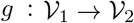 be a one-to-one mapping between the vertices of the two networks. For an edge 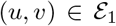, we call 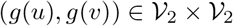 to be a translation of (*u, v*) from 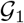 to 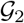. Then, the metric EC, which measures the percentage of the edge-translation of edges in 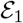 that is an actual edge in 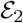, is mathematically described as

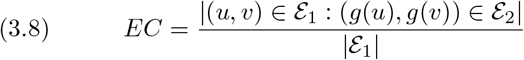

Since the PPI connections are (weakly) preserved during evolution, higher EC values imply better protein pairings.

#### Largest Common Connected Subgraph (LCCS)

Given two graphs, 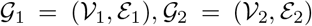, with 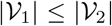, and a one-to-one mapping 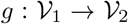, let *E* denote the set of edges of 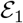 that map to edges in 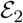 using *g*. Then 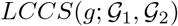 denotes the number of edges in the largest connected component of *E*. Like EC, LCCS also measures the number of edges that are preserved across the species by the mapping *g*.

#### Average Functional Similarity

The two previous metrics calculate how well the protein mapping preserves the structural features of the PPI networks. Since important biological functional pathways are also broadly preserved during evolution, we also expect a good mapping to strongly preserve protein function across species. To evaluate this, we employ a widely used taxonomy of protein functions called the Gene Ontology (GO) [4]. The GO is a hierarchical and species-agnostic Directed Acyclic Graph (DAG) representation of protein functional labels (i.e. GO-terms) composed of three domains (Molecular Function (MF), Biological Process (BP) and Cellular Component (CC)), each with a separate root. The GO taxonomy is hierarchical; so the GO-Terms closer to the roots represent more general protein functions, which becomes increasingly specialized as we move farther away from the roots. We depart from [8] to match recommended best practice in the field, and enforce some specificity of protein function by including only the labels whose shortest distance from a root is ≥ 5. We associate all nodes with their known GO terms as collected in the GO database [4] where we note that proteins are typical annotated with multiple GO-terms. So, for each protein mapping (*p, g*(*p*)) we use Schlicker’s similarity (i.e. *s_c_*(*p, g*(*p*)), *c* ∈ {MF, BP, CC}) [15, 8] to find the similarity of GO-Terms of *p* with that of *g*(*p*). So, the Average Functional Similarity (*AFS_c_*) is computed as

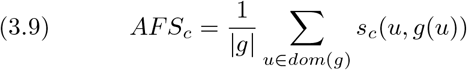

Where |*g*| represents the size of one-to-one protein mappings in *g* (If the alignment is global, 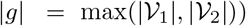. Mappings with higher AFS based on the known functional labels are preferred.

##### 3.2.1 Approximation Loss and Matching Similarity

In order to see how the IsoRank approximations compare with the exact IsoRank mappings according to these measures, we performed a similar analysis to that done in Section 3.1. For a given pair of species (*A, B*), *A* ≠ *B*, let *R*(*α; A, B*) be the true IsoRank similarity matrix, and let *R*^0^(*α; A, B*) and *R*^1^(*α; A, B*) be the approximations described in (2.5) and (2.7) respectively, for a given *α*-value. After using the greedy matching algorithm on *R*(*α; A, B*), *R*^0^(*α; A, B*) and *R*^1^(*α; A, B*), let *M*(*α; A, B*), *M*^0^(*α; A, B*) and *M*_1_ (*α; A, B*) be the corresponding list of 2000 top one-to-one cross-species protein matches for the species pair (*A, B*). All these results are generated for *α* ∈ {0.2, 0.4, 0.6, 0.8} and *A, B* ∈ *S*.

Figure 1(A) shows the percentage similarity of mappings *M*^0^ and *M*_1_ with respect to the true mapping *M* for different *α*-values. We find that percentage similarities are broadly stable with *α* and in all species pairs, *M*_1_ mapping is closer to *M* than *M*^0^. The disparity between M^0^ and *M*_1_ is most pronounced in the fly-yeast pair, where the difference is > 0.25 for all *α*. However, for pairs where one of the organisms is human, there is relatively minor difference between the mapping produced by the two approximations, and the mappings themselves are closer to the true mapping.

**Figure 1:**
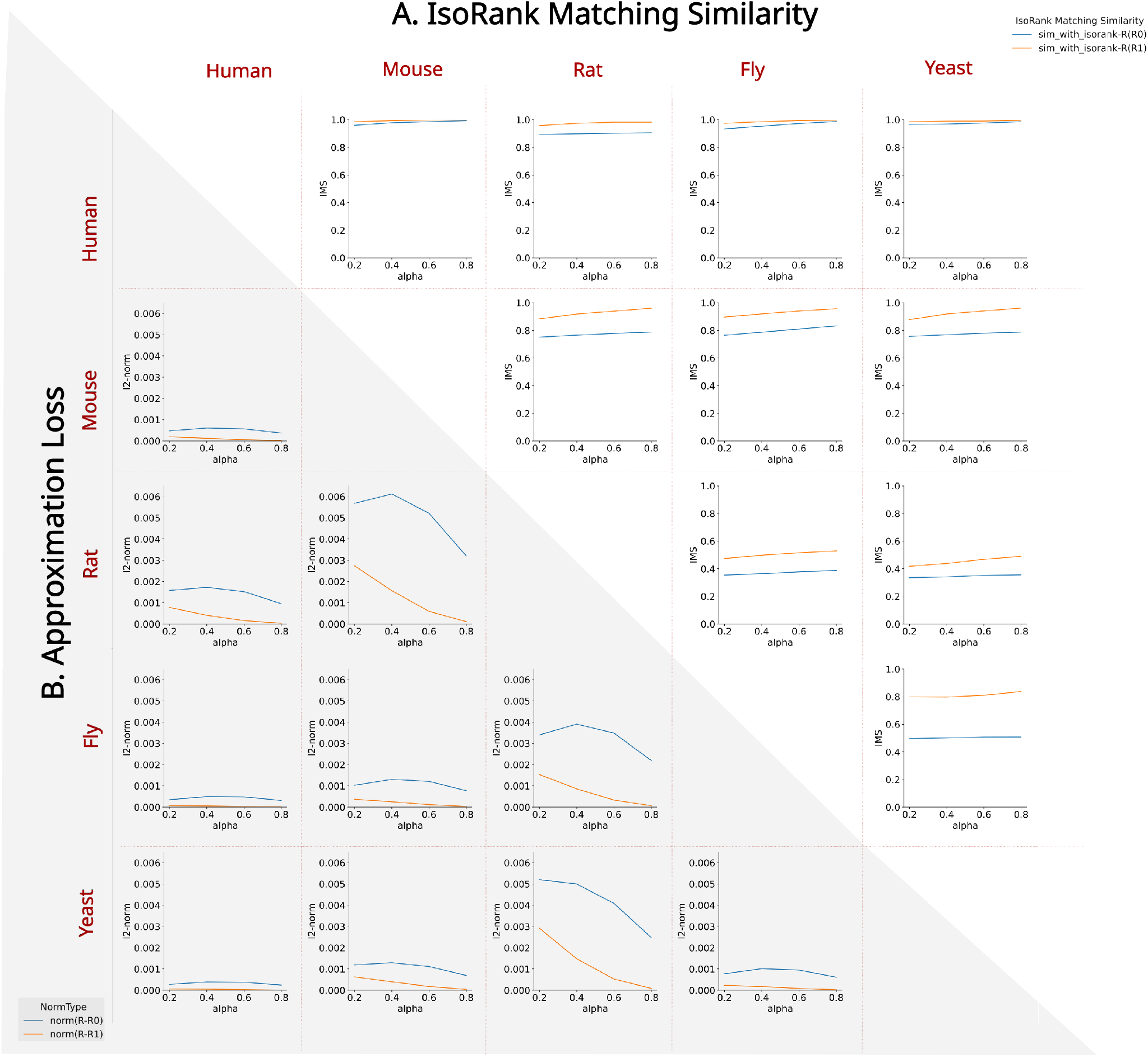
Plots of A) IsoRank Matching Scores and B) Approximation loss for different *α*-values obtained from the true IsoRank similarity matrix *R* and its two approximations *R*^0^ and *R*^1^ for all pairwise combinations of organisms: Human, Mouse, Rat, Fly and Yeast.

Figure 1(B) shows the plot of L2-norms ||*R*^0^(*α; A, B*) – *R*(*α; A, B*)|| and ||*R*^1^(*α; A, B*) – *R*(*α; A, B*)|. Unlike in 1(A), we see that the L2 loss is more sensitive to *α*. For all species pairs, the approximation loss is usually the highest when *α* ∈ (0.4, 0.6) and the lowest when *α* → 1. In all scenarios, the approximation loss is below 0.0075.

##### 3.2.2 EC and LCCS results

Figure 2(A) shows the variation of LCCS scores for IsoRank and its approximations with respect to *α* values for all distinct species-pairs of *S*. Even though the true IsoRank mapping always produces the best LCCS scores in all situations tested, the results obtained from the *R*^1^ approximation described in (2.7) are rarely much worse. In some species pairs (for example Fly-Human, Mouse-Fly and Rat-Fly) even the *R*^0^ results closely match the LCCS scores obtained from true IsoRank. The same pattern is observed when we used the EC scores to compare the approximations (Figure 2(B)), though we see an exception in the Baker’s Yeast-Rat alignment, where the gaps between the EC scores of true IsoRank and its approximations are ≥ 0.2.

**Figure 2:**
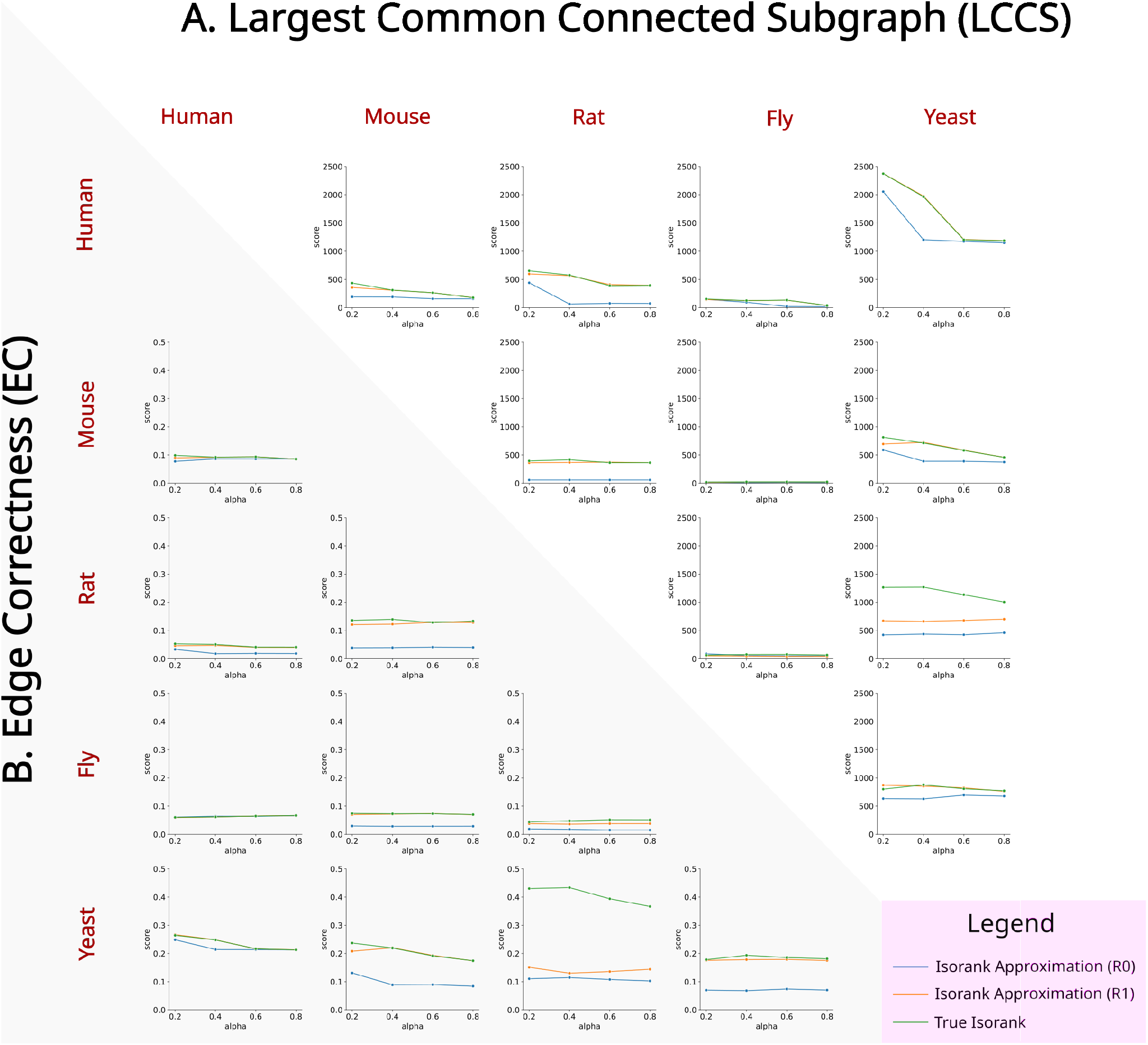
Plots of A) EC and B) LCCS scores for different *α*-values obtained from the true IsoRank similarity matrix *R* and its two approximations, *R*_0_ and *R*_1_ for all pairwise combinations of organisms: Human, Rat, Mouse, Fly and Yeast. The results are computed using the 2000 top-scoring matches.

##### 3.2.3 Functional Similarity Results

We used the Average Functional Similarity (AFS) measure, described above, to measure how functionally similar the protein-pairs mapped by IsoRank and its approximations are. The AFS results are provided in Table 8. Since the AFS result was not sensitive to *α*-values (fixing GO-category, species-pairs and type of IsoRank, the expected s.d. obtained by setting *α* = {0.2, 0.4, 0.6, 0.8} was ≈ 0.0015), we report *α* = 0.6 in Table 8 and present the obtained AFS scores across all GO-categories and species-pairs.

**Table 8:** Average Functional Similarity (AFS) results for top 2000 matching obtained from two versions of approximate IsoRank (***R***^0^ and ***R***^1^) and true IsoRank (***R***) matrices. for MF, BP and CC GO hierarchies, with *α* = 0.6.

| org1 | org2 | AFS($R^0$) | AFS($R^1$) | AFS($R$) |
| --- | --- | --- | --- | --- |
| Molecular Function |  |  |  |  |
| fly | bakers | 0.513322 | 0.515298 | **0.516975** |
| fly | mouse | **0.574128** | 0.573517 | 0.570199 |
| fly | rat | **0.495021** | 0.429881 | 0.391325 |
| human | bakers | **0.595207** | 0.594926 | 0.595034 |
| human | fly | **0.630536** | 0.627544 | 0.627822 |
| human | mouse | **0.661401** | 0.659792 | 0.659436 |
| human | rat | **0.622408** | 0.621161 | 0.621160 |
| mouse | bakers | 0.515291 | **0.523203** | 0.521085 |
| mouse | rat | **0.613179** | 0.600434 | 0.600229 |
| rat | bakers | **0.477499** | 0.426828 | 0.360398 |
| Biological Process |  |  |  |  |
| fly | bakers | 0.260714 | 0.266032 | **0.267592** |
| fly | mouse | 0.267876 | **0.269937** | 0.269391 |
| fly | rat | **0.190414** | 0.170344 | 0.154421 |
| human | bakers | **0.361614** | 0.361569 | 0.361258 |
| human | fly | **0.334539** | 0.331662 | 0.331553 |
| human | mouse | **0.347836** | 0.345505 | 0.345371 |
| human | rat | **0.308425** | 0.307742 | 0.307717 |
| mouse | bakers | 0.264904 | 0.273303 | **0.274346** |
| mouse | rat | **0.297277** | 0.292720 | 0.291989 |
| rat | bakers | **0.215697** | 0.188325 | 0.159221 |
| Cellular Component |  |  |  |  |
| fly | bakers | 0.538923 | 0.546564 | **0.550457** |
| fly | mouse | 0.579008 | **0.586186** | 0.583203 |
| fly | rat | **0.428872** | 0.399232 | 0.368953 |
| human | bakers | 0.502686 | **0.502984** | 0.502560 |
| human | fly | **0.585266** | 0.581585 | 0.581809 |
| human | mouse | **0.575433** | 0.573686 | 0.573678 |
| human | rat | **0.464256** | 0.464092 | 0.464261 |
| mouse | bakers | 0.525856 | **0.529648** | 0.529438 |
| mouse | rat | **0.536556** | 0.530721 | 0.529979 |
| rat | bakers | **0.409277** | 0.376957 | 0.344751 |

The results are in stark contrast to the results obtained from network-sensitive measures, discussed in Section 3.2.2. Unlike EC and LCCS outputs, the mapping obtained from the *R*^0^ approximation almost always gave the highest AFS scores across all GO categories (though in most case by a slight margin). So, for the measure the biologists care most about, it seems that the approximate IsoRank is superior to exact IsoRank. Even in cases where it didn’t produce the highest-scoring one-to-one mapping, it closely followed the best scoring algorithm.

## 4 Discussion

### 4.1 Synthetic Networks

As shown from Table 1 through 6, for both ER and BA random graphs, when “good” ***E*** is used, the accuracy for all three algorithms is 100%, i.e., all the algorithms can recovery the network alignments perfectly, and the errors, measured in both norms, are very small. However, the original IsoRank 2.2 is much slower than the two approximation algorithms. Approximate IsoRank 2.5 is about 7.6-14.3 times faster, and approximate IsoRank 2.7 is about 4.7-6.7 times faster. Since they all recover the perfect alignment, when ***E*** contains accurate information about the alignment, we suggest using approximate IsoRank 2.5. When “in-between” ***E*** is used, although the error between the original IsoRank and approximations are still small, the accuracy for all three algorithms decreases. As we can see, Approximate IsoRank 2.5 is the fastest among the three, but its accuracy is slightly worse than the original IsoRank 2.2. Approximate IsoRank 2.7 is slightly slower than approximate IsoRank 2.5 but achieves roughly the same accuracy as the original IsoRank 2.2. But approximate IsoRank 2.7 is still much faster than original IsoRank 2.2 and, therefore, balancing the accuracy and CPU time, we would suggest using approximate IsoRank 2.7 in this case. Finally, when ***E*** is “bad”, we observe the same results as the case ***E*** is “in-between”. In particular, approximate IsoRank 2.5 is the fastest, but its accuracy is the worst (but not by much, still close to the accuracy of the original IsoRank 2.2). Approximate IsoRank 2.7 seems to achieve the best balance between the accuracy and CPU time, and we would suggest using it in this case as well. However, if the CPU time is the priority, then we recommend approximation IsoRank 2.5 since it is the fastest. Finally, from the numerical experiments, we observe that both our approximation algorithms achieve optimal computational complexity 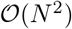, where 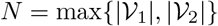, as expected since the synthetic networks considered here are sparse (namely 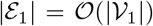) and 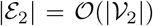 and 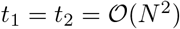 in our implementation.

### 4.2 Biological Networks

Interestingly, we found the *R*^0^ approximation to IsoRank to be more robust and produce better predictions of protein functional roles across the board than even exact IsoRank. Since inferring the function of unknown proteins across species is one of the main applications of the Global Alignment of Biological Networks, perhaps given the fact that the networks aren’t actually isomorphic plus we have an imperfect sample of the true edges, means that somehow our approximation to IsoRank is more robust to noise, and therefore works better than producing the best isomorphism as measured by purely graph-theoretic measures. This requires further investigation.

## Code and Data availability and Funding Acknowledgements

Temporarily removed for anonymous author submission. All code and data will be made available on GitHub. Thanks to NSF (grant number removed).

The Gene Ontology files are taken from the [official GO website](http://geneontology.org/) (OBO data version: releases/2022-12-04, GAF version: 2.2)

The biological networks used in the experiments can be downloaded from the [IntAct FTP link](https://www.ebi.ac.uk/intact/download/ftp).

